# Paclitaxel Alleviates Pulmonary Arterial Hypertension by Modulating Macrophage M2a Polarization via the NF-κB Pathway

**DOI:** 10.1101/2025.08.15.670469

**Authors:** Yong Wu, Ling Gu, Hanmin Liu, Li Wang, Qiwei Li, Lu Zhu

**Affiliations:** Mianyang Central Hospital, School of Medicine, University of Electronic Science and Technology of China, Mianyang, Sichuan, China; Key Laboratory of Birth Defects and Related Diseases of Women and Children (Sichuan University), Ministry of Education, West China Second University Hospital, Sichuan University, Chengdu, Sichuan, China; Southwest Medical University, Luzhou, Sichuan Province, China

**Keywords:** pulmonary arterial hypertension, M2a macrophage, paclitaxel, NF-κB Signaling pathway

## Abstract

**Background:** Pulmonary arterial hypertension (PAH) is primarily associated with vascular remodeling, in which macrophage polarization is considered to play a major role. Paclitaxel(PTX) has been shown to modulate macrophage polarization in tumors. We investigated whether PTX could alleviate PAH by altering macrophage polarization, particularly M2a macrophages.

**Methods:** To establish PAH, rats underwent left pneumonectomy followed by monocrotaline (MCT) injection. In vitro, THP-1 cells were differentiated into macrophages using PMA and polarized with LPS+IFN-γ (M1), IL-4 (M2a), or IL-10 (M2c), followed by PTX intervention. Hemodynamic parameters (mPAP) and right ventricular hypertrophy index (RVHI) were measured. Pathological changes were assessed via HE staining. Single-cell RNA sequencing analyzed macrophage subsets in lung tissues. Macrophage polarization and NF-κB pathway activity were evaluated using immunohistochemistry, qPCR, cytokine profiling and Western blot.

**Results:** In a severe PAH model induced by left pneumonectomy plus monocrotaline, pulmonary interstitial macrophages initially exhibited a predominant M1-polarized phenotype in the early stage, which transitioned to an M2a-polarized phenotype in the later stage and eventually stabilized. Low-dose PTX alleviated severe PAH by reducing perivascular macrophage infiltration. Cytological studies confirmed that PTX inhibited M2a polarization via suppression of NF-κB, thereby attenuating its proliferative effects on pulmonary arterial smooth muscle cells (PASMCs).

**Conclusion:** M2a macrophages play a crucial role in PAH progression. PTX may alleviate PAH by suppressing M2a polarization via the NF-κB pathway.

## INTRODUCTION

Pulmonary arterial hypertension (PAH), a fatal subgroup of pulmonary hypertension(PH), is characterized by vascular remodeling, inflammation, and elevated pulmonary vascular resistance[1]. Current therapies primarily target vasodilation but fail to reverse pathological remodeling [2–5]. Emerging evidence implicates macrophage-driven inflammatory microenvironments in PAH progression [6]. Macrophages exhibit plasticity, polarizing into pro-inflammatory M1 or anti-inflammatory M2 subtypes, with M2 macrophages promoting pulmonary artery smooth muscle cells (PASMC) proliferation and vascular remodeling [7–8]. Further research has classified M2 macrophages into M2a, M2b, M2c, and M2d subtypes[9,10]. But the roles of these M2 macrophage subtypes in PAH is still not well understood.

Paclitaxel (PTX) is a chemotherapeutic agent that reprograms tumor-associated macrophages (TAMs)M2 into the M1 phenotype via the TLR4-NF-κB signaling pathway [11]. However, whether it exerts the same effect in PAH remains unclear. Therefore, we investigated the effects of PTX on the polarization of pulmonary interstitial macrophages in PAH and explored the underlying mechan isms of PTX on M2 subtype macrophages.

## METHODS

### Experimental PAH models

The PAH rat model was established as follows: After one week of acclimatization, Sprague-Dawley (SD) rats underwent left pneumonectomy. One week post-surgery, a single dose of monocrotaline (MCT, 60 mg/kg) was administered via subcutaneous injection in the neck region. At the 4th week (4W) and 6th week (6W) post-operation, paclitaxel-loaded nanoparticles (2 mg/kg) were intravenously injected via the tail vein. The detailed experimental protocol is illustrated as previously described.[12,13] (Fig S1 A in Supporting Information).

### Materials and Reagents

Reagents used for animal model establishment and associated experiments are listed in Supplementary Material S1.

### Hemodynamic measurements and Collection of heart and lung tissues

Right ventricular direct vision manometry was used to measure 3 to 6 systolic and diastolic pressures in each rat. Then, cardiac and lung tissue specimens were collected for the next step of the experiment. (see Supporting Information S1).

### Single-Cell Sequencing of Whole Lung Tissue

Following cardiac pressure measurement and pulmonary vasculature saline perfusion, one lobe from each of the left and right lungs was excised. The tissues were minced and digested with dispase and collagen mixed enzymatic solution using a Miltenyi Biotec tissue homogenizer (Miltenyi Biotec, Germany) to generate a single-cell suspension. Cell suspensions meeting the following criteria were processed: >80% viability (assessed by trypan blue exclusion), a concentration of 700–1,200 cells/μL, a diameter range of 5–30 μm, and a total yield of >100,000 cells. Sequencing was carried out employing the Illumina NovaSeq PE150 paired-end sequencing strategy.

### Histopathology and Immunostaining

Lung tissues were fixed, paraffin-embedded, and stained with HE. Macrophage markers (CD68, CD86, CD206), α-SMA, TLR4, and NF-κB were analyzed via immunohistochemistry.

### Cell Culture

THP-1 cells were differentiated into macrophages with PMA (100 ng/ml) and polarized with LPS+IFN-γ (M1), IL-4 (M2a), or IL-10 (M2c) (Supplementary S1). PTX (10 μM), TLR4 inhibitor (VGX-1027), or NF-κB inhibitor (EVP4593) were applied for 24 h. The effects of supernatants from different macrophage subtypes on the proliferation capacity of human smooth muscle cells were evaluated.. Human pulmonary artery smooth muscle cells (PASMC) are from Mei sen CTCC.Detailed intervention methods can be found in Supporting Information S1.

### Molecular Analysis

Cytokine levels (TNF-α, IL-4, IL-10) were quantified using Luminex assays. Gene expression of CD68, CD86, Mrc1, IL-4 and protein levels (TLR4, NF-κB, CD206, TNF-α) were assessed via RT-qPCR and Western blot.The sequence of primers is shown in Supporting Information Table S1.

### Statistical Analysis

Statistical analyses were performed using one-way ANOVA (Tukey’s multiple comparisons test, Dunnett’s multiple comparisons test and Dunn’s multiple comparisons test), two-way ANOVA (Sidak’s multiple comparisons test) and t-test (Prism 9; GraphPad Software, La Jolla, CA, USA).Differences were considered significant at p <0.05.

## RESULTS

### M2a promote the development of PAH in rats

The PAH rats exhibited elevated mPAP and Rightt ventricular hypertrophy index (RVHI) at 4 weeks post-surgery(Fig S1 B,C, in the Supporting Information). Followed by the development of severe PAH accompanied by significant vascular remodeling at 6 weeks postoperatively (Fig S2 G, H in Supporting Information).

Subsequently, we employed single-cell RNA sequencing to analyze pulmonary tissue macrophages. The enrichment of whole lung tissue cells resulted in the classification into 29 distinct cell clusters (Fig 1A). Comparative analysis across different samples revealed a significant increase in the proportion of cluster 8, identified as pulmonary interstitial giant cells, at 4 weeks and 6 weeks post-operation (Fig 1B). Our findings revealed that the genes C1qa, C1qb, and C1qc were specifically expressed in pulmonary interstitial macrophages (Fig 1 C). Moreover, among C1qc+C1qa+C1qb+CD68+ pulmonary interstitial macrophages, C1qc+C1qa+C1qb+CD68+CD86+Fcgr1a+ M1-like macrophages were highly expressed in the early stage of PAH (4 weeks post-surgery). In contrast, during the late stage of PAH (6 weeks post-surgery), C1qc+C1qa+C1qb+CD68+Mrc1 (CD206)+ M2a-like macrophages exhibited significantly higher expression compared to M1 macrophages.((Figure1 D)

**Fig. 1.**
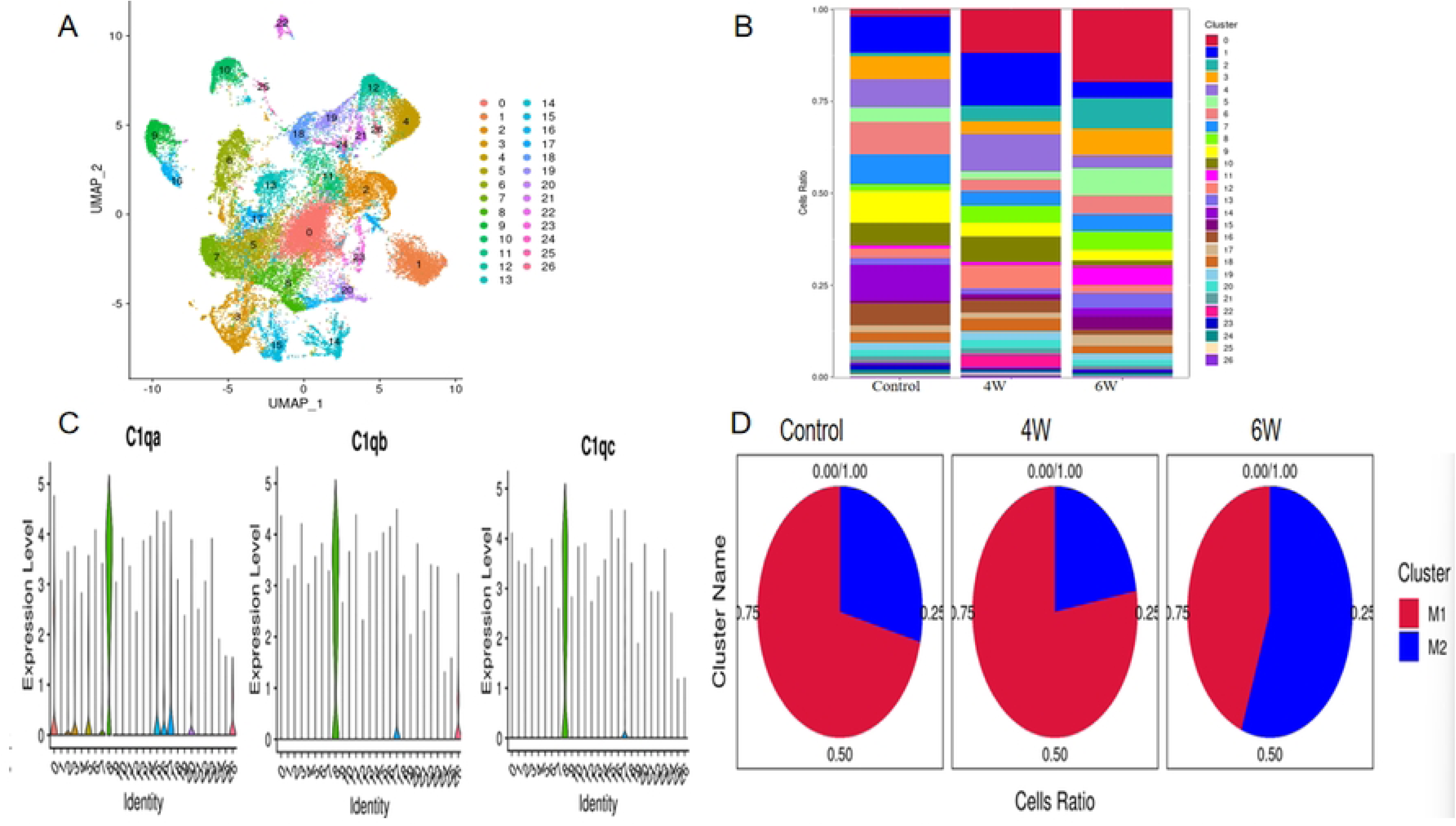
Single-cell sequencing results related to whole lung tissue. A. The lung tissue cells were enriched based on different markers and classified into 29 distinct cell clusters, ranging from cluster 0 to cluster 28. B. A comparison of the proportions of different cell types between the PAH groups (4W, 6W) and the control group at different time points revealed that the proportion of pulmonary interstitial macrophages in the 8 groups was significantly higher in the PAH group than in the control group. C. Single-cell sequencing revealed that the expression of the C1qa, C1qb, C1qc gene in group 8 pulmonary interstitial macrophages was significantly higher than that in other cell populations. D. In the control group of single-cell sequencing, the pulmonary interstitial macrophage population was predominantly composed of M1 macrophages (more than 50% but less than 75%),. In the early stage of PAH group (4W, post-surgery), M1-like macrophage was significantly increased (>75%). In the late stage of PAH group (6W, post-surgery), M2a macrophage polarization was markedly elevated (>50%)

Further immunohistochemical analysis confirmed that CD68+ macrophages exhibited the most pronounced perivascular aggregation at 4 weeks post-operation(4W), with concurrent increases in both CD86+ M1 and CD206+ M2a subtypes (Fig 2A-E). The macrophages displayed cellular hypertrophy, with the majority of interstitial macrophages exhibiting a foamy morphology. Quantitative analysis revealed that the number of CD86+ cells was significantly higher than CD206+ cells at 4 weeks, whereas this ratio was reversed at 6 weeks when CD206+ cells outnumbered CD86+ cells. Notably, CD68+ macrophages maintained substantial perivascular accumulation during both 6-week and 8-week PAH stages (Fig 2 A-E).

**Fig 2.**
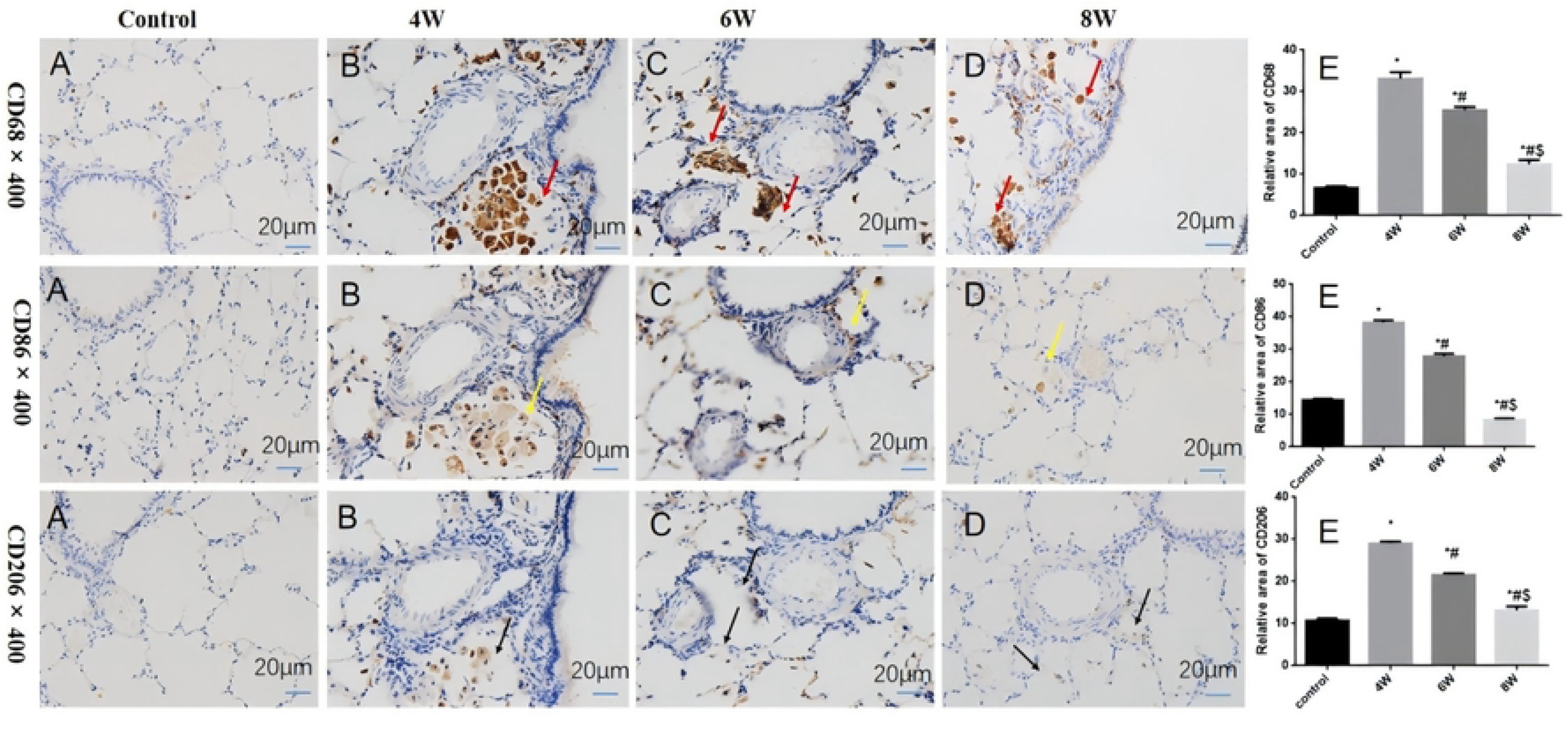
The expression of CD68, CD86, and CD206 proteins in the perivascular regions of rat pulmonary arteries in adjacent tissue sections. A. The control group showed that the protein expressions of CD68, CD86 and CD206 were not obvious; B. 4 weeks post-surgery group showed CD68+ macrophages, CD86+ M1-like macrophages, and CD206+ M2a-like macrophages aggregated around the pulmonary arteries, with a significant increase in macrophage count. The protein expression levels were markedly higher than those in the control group, and the expression of CD68+ and CD86+ proteins was higher than that of CD206+. C. 6 weeks post-surgery group showed CD68+ macrophages persist around the pulmonary arteries, CD86+ macrophages are reduced, and CD206+ macrophages decrease compared to 4W but remain significantly present D. 8 weeks post-surgery group showed CD68+ macrophages persist around the pulmonary arteries, CD86+ macrophages continue to decrease, and CD206+ macrophages persist and are more abundant compared to the control group. E. Quantification of relative protein expression levels of CD68, CD86, and CD206 across various stages of pulmonary arterial hypertension (PAH) using ImageJ software. The observation under the microscope showed that the background was purple-blue, while the positive proteins were stained brown-yellow or yellow. 4W:4 weeks after left pneumonectomy. 6W: 6 weeks after left pneumonectomy 8W:8 weeks after left pneumonectomy. *P<0.05, compared with the Control group; # P<0.05, compared with 4W group; $ P<0.05, Compared with 6W group,

Comparative evaluation demonstrated a marked reduction in CD86+ macrophage counts at 6 and 8 weeks compared to the 4-week timepoint. Although CD206+ M2a-like macrophage aggregation around vasculature decreased at 6 and 8 weeks relative to 4 weeks, their numbers remained significantly elevated compared to control groups (Fig 2 A-E).

Meanwhile, we found that TLR4 and NF-κB proteins exhibited significant expression in perivascular macrophages at 4, 6, and 8 weeks (4W, 6W, 8W) in PAH, with peak NF-κB expression detected in macrophages at 4 weeks after operation(Fig S3 A-E in Supporting Information).

### Paclitaxel attenuates PAH and Decreases Perivascular M2a Macrophages

After early intervention (4W) with paclitaxel for two weeks, no significant difference was observed in mPAP compared to 6W PAH rats, while RVHI was reduced (Fig 3 C,G). In contrast, after late intervention (6W) with paclitaxel for two weeks, both mPAP and RVHI were decreased compared to PAH rats at the same time poin (Fig 3 D,H).

**Fig 3.**
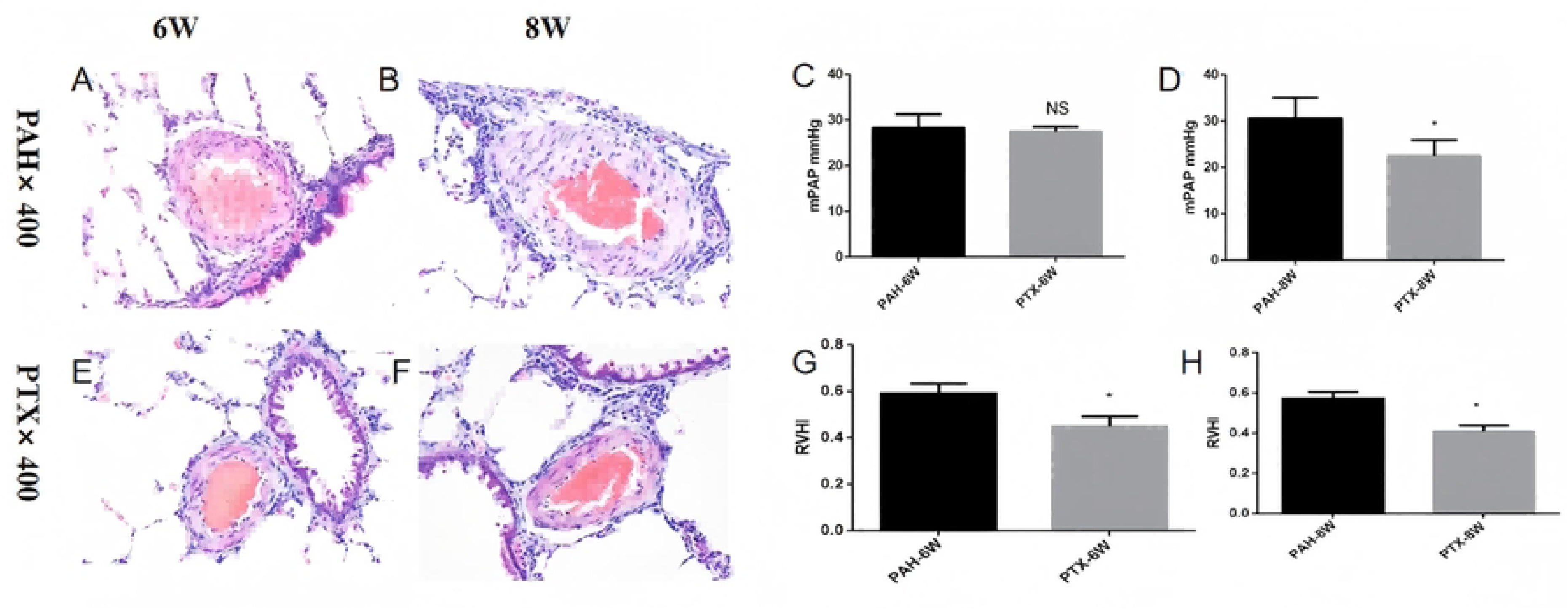
Comparison of mPAP and RVHI between PAH rats after 2-week PTX intervention and PAH rats of the same period. A–B. Haematoxylin and eosin (H&E) staining demonstrated significant thickening of the pulmonary arterial smooth muscle layer at 6 weeks (6W) and 8 weeks (8W) post-surgery. C–D. Haemodynamic assessment revealed that mean pulmonary arterial pressure (mPAP) remained unchanged at 6W but was significantly reduced at 8W following paclitaxel (PTX) treatment (P < 0.05 vs. baseline). E–F. Comparative histological analysis showed that, relative to untreated controls, PTX intervention did not alter pulmonary arterial smooth muscle thickening at 6W but significantly attenuated it by 8W (P < 0.05). G–H. The right ventricular hypertrophy index (RVHI) was significantly lower in PTX-treated animals at both 6W and 8W compared to the non-intervention group (P < 0.05). * vs. PAH group p<0.05

At the same time, in the later stage of PAH (6 weeks), a two-week intervention with paclitaxel can reduce the perivascular accumulation of CD206+ M2a macrophages. (Fig 4A). Use Imagine J statistic revealed that early intervention with paclitaxel at 4 weeks (4W) in PAH rats for 2 weeks resulted in a reduction of CD68+ macrophages around the pulmonary arteries compared to age-matched PAH controls, while the expression of CD86+ M1 and CD206+ M2a macrophages showed no significant difference (Fig 4B). In contrast, after 2 weeks of paclitaxel intervention during the late stage (6W) of PAH, both CD68+ macrophages and CD206+ M2a macrophages in the pulmonary arterial perivascular area were reduced compared with age-matched PAH rats, while CD86+ M1 macrophages showed no significant chang (Fig 4C).

**Fig 4.**
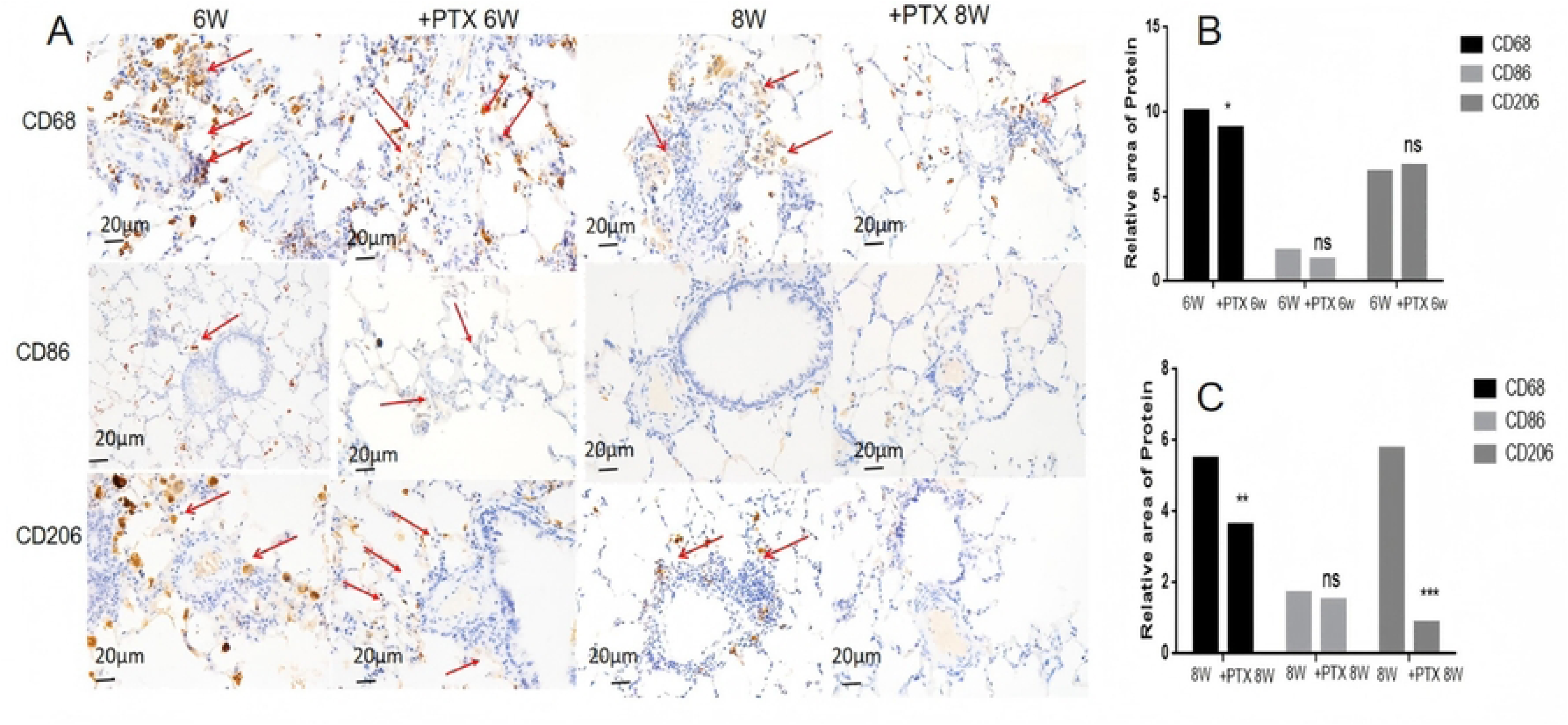
After PTX intervention, the expression differences of macrophage-related proteins (CD68, CD86, CD206) in lung tissues compared with pulmonary arterial hypertension rats of the same period. A. Representative immunohistochemical images of different stages of PAH using CD68, CD86, and CD206 as markers The observation under the microscope showed that the background was purple-blue, while the positive proteins were stained brown-yellow or yellow.. The red arrows indicate macrophages under different markers B. Statistical comparisons were conducted using one-way ANOVA followed by Tukey’s multiple comparisons test versus 6W or +PTX 6W and are represented above each plot. *p <0.05, **p <0.01.***p <0.001. C.Statistical comparisons were conducted using one-way ANOVA followed by Tukey’s multiple comparisons test versus 8W or +PTX 8W and are represented above each plot. *p <0.05, **p <0.01.***p <0.001. 6W: 6 weeks after left pneumonectomy 8W:8 weeks after left pneumonectomy. +PTX 6W: PAH rats were treated with PTX for 2 wee ks starting at 4 weeks after surgery. +PTX 8W: PAH rats were treated with PTX for 2 wee ks starting at 6 weeks after surgery.

### M2a Stimulates PASMC Proliferation, While Paclitaxel Alleviates PAH by Inhibiting M2a Polarization via Suppression of the NF-κB Pathway

In vitro cell experiments, we co-cultured PASMCs with conditioned media from different macrophage subtypes for 72 hours and found that M2a-conditioned media consistently and strongly stimulated PASMC proliferation (Fig S4 in Supporting Information).

Previous immunohistochemical studies have confirmed the high expression of TLR4 and NF-κB in macrophages of PAH. In vitro experiments we found that the addition of PTX downregulated NF-κB gene expression, while it exhibited no significant effect on TLR4 gene expression. Furthermore, the NF-κB inhibitor EVP4593 significantly suppressed NF-κB gene expression (Fig 5A).

**Fig 5.**
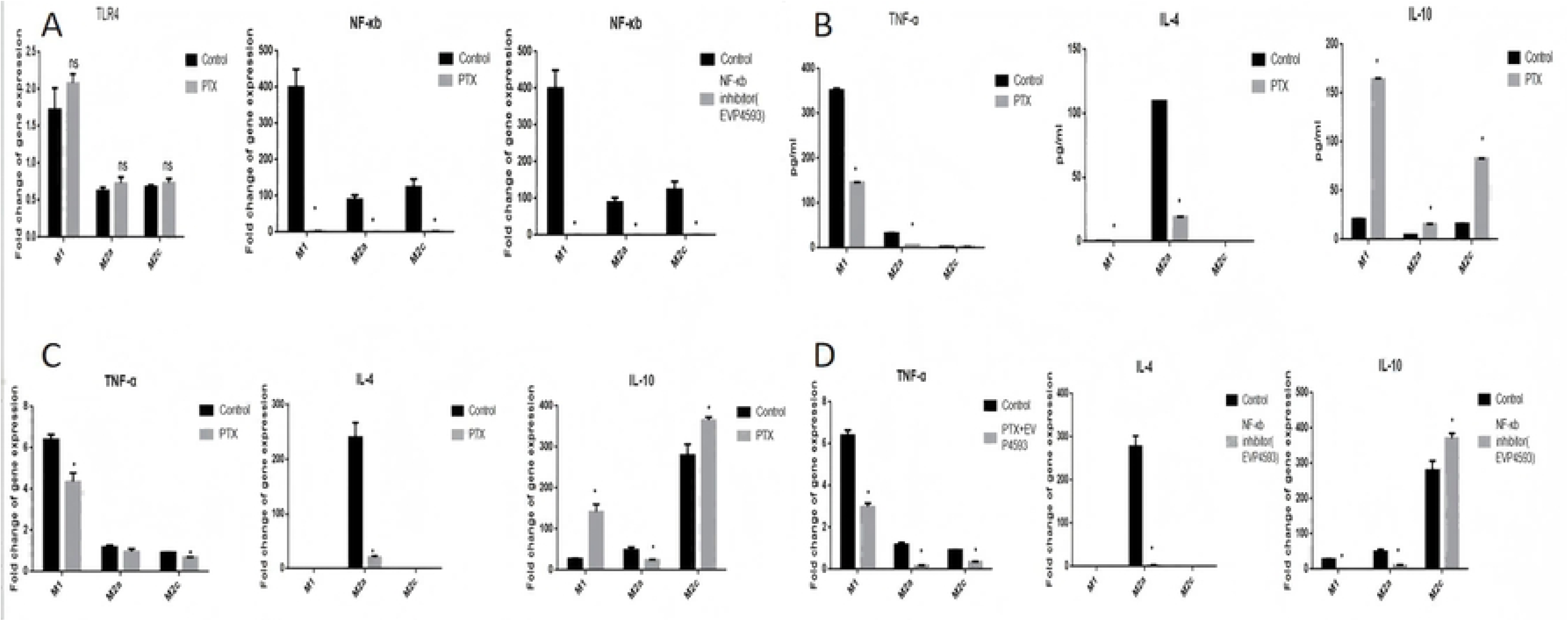
Differences in macrophage-related cytokines and gene expression after PTX intervention. A. Effects of PTX and NF-κB inhibitor EVP4593 on TLR4/NF-κB expression in macrophage subsets (M1, M2a, M2c), Both PTX and the EVP4593 significantly suppressed NF-κB gene expression. B.PTX intervention significantly reduced the levels of TNF-α and IL-4 cytokines in macrophages, increasing the level of IL-10. C.After PTX intervention, the gene expression of TNF-α and IL-4 significantly decreased, while the expression of IL-10 increased. D. After intervention with the NF-κB inhibitor EVP4593, the gene expression of TNF-α and IL-4 decreased, while the gene expression of IL-10 increased. Results are presented as means±SD from five to 10 samples. Statistical comparisons of multiple groups were conducted using ANOVA followed by Tukey’s multiple comparisons test. *p <0.05.

Simultaneously, the liquid chip assay revealed that PTX intervention significantly reduced the levels of TNF-α and IL-4 cytokines in macrophages, while increasing the level of IL-10 (Fig 5B). A corresponding trend was also observed at the gene expression level (Fig 5C). Furthermore, after the addition of the NF-κB inhibitor EVP4593, the expression patterns of the relevant genes were similar to those observed following PTX intervention (Fig 5D).

Further protein analysis of M2a cells revealed that PTX intervention downregulated the expression of CD206, an M2a-associated protein, accompanied by reduced expression of TLR4 and NF-κB proteins (Fig 6A-B). Additionally, the NF-κB inhibitor EVP4593 further decreased CD206 protein expression (Fig 6 D).

**Fig 6.**
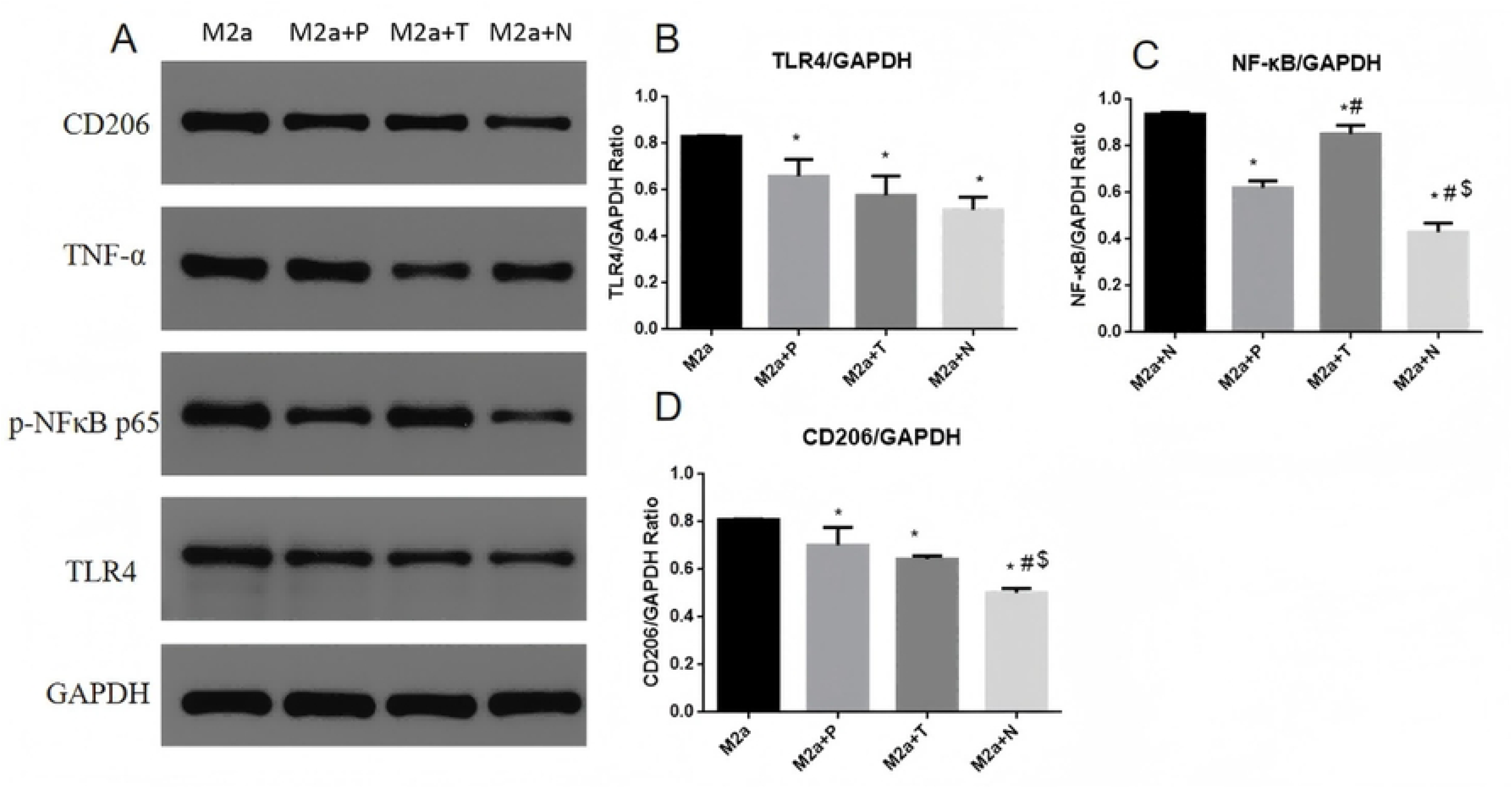
Differences in macrophage-related protein expression after PTX intervention. A. Protein expression profiles of M2a cells following treatment with different interventions (PTX, TLR4, and NF-κB antagonists). B. Following treatment with various interventions(PTX, TLR4, and NF-κB antagonists), TLR4 protein expression in M2a macrophages was consistently downregulated C. Effects of different interventions(PTX, TLR4, and NF-κB antagonists). on CD206 protein expression in M2a macrophages: All treatments resulted in downregulation of CD206 protein expression. * vs M2a P<0.05. M2a+P: PTX added to the M2a group; M2a+T: TLR4 inhibitor VGX-1027 added to the M2a group; M2a+N: NF-κB inhibitor EVP4593 added to the M2a group.

## DISCUSSION

In this study, we report that the M2 macrophage subtype M2a may be a key immune cell responsible for inducing PAH. Furthermore, we demonstrate that paclitaxel likely alleviates severe PAH by suppressing M2a macrophages via the NF-κB pathway.

Polyarterial arterial hypertension (PAH) is a form of pulmonary hypertension characterized by elevated pulmonary artery pressure resulting from increased pulmonary vascular resistance [14]. Without treatment, PAH typically progresses to right ventricular failure and ultimately leads to death. Current understanding attributes its vascular pathology to medial and intimal remodeling, plexiform lesions, and fragmentation of the elastic layer [15]. This pathological phenotype can be induced by various environmental stressors and damaging stimuli, including inflammatory responses, DNA damage, and epigenetic dysregulation. Following endothelial cell injury, pro-inflammatory cells are recruited, promoting the infiltration of lymphocytes, macrophages, and dendritic cells into the lungs. This perpetuates local inflammation and autoimmune responses, ultimately contributing to a distinct pulmonary smooth muscle cell phenotype. These cells acquire migratory capabilities, leading to vascular wall thickening and muscularization [16].

The early study observed a gradual increase in CD68+ macrophages over three weeks in a SU + hypoxia-induced PAH model using athymic rats[17]. This finding aligns with our experimental results in the early stage of PAH (4W). However, in the late stage of PAH (6W), although CD68+ macrophage expression remained significantly elevated, it did not exhibit a continuous upward trend. This suggests that macrophages play a sustained role in PAH progression, stabilizing at a certain level in the later stages to exert their effects. Meanwhile, we observed that CD86+ M1-like macrophages predominated in the early stage of PAH (4W), while CD206+ M2-like macrophages were dominant in the late stage (6W), further confirming the dynamic polarization of macrophages during PAH progression.

Further studies have revealed that M2 macrophages play a significant role in the progression of PAH[18,19,20,21]. Based on the specific stimuli received and the resulting phenotypic and functional characteristics, M2 macrophages can be further classified into four distinct subtypes: M2a, M2b, M2c, and M2d.[22,23,24]. M2a macrophages exhibit limited capacity to induce T cell activation and proliferation, thereby potentially facilitating tumor growth[25,26].These macrophages are characterized by elevated expression of the cell surface markers CD206, CD209, and Dectin-1[27]. CD206 is commonly used as a representative marker for M2a macrophages.[10,28,29]. In our study, single-cell sequencing and immunohistochemistry in PAH rats revealed elevated expression of CD206-associated genes and proteins, further confirming the active role of CD206+ M2a-like macrophages in PAH progression. M2b macrophages can be induced by combined exposure to immune complexes (IC) and Toll-like receptor (TLR) agonists or interleukin-1 receptor (IL-1R) agonists, and they express high levels of CCL1 and tumor necrosis factor superfamily member 14 (TNFSF14)[30]. These M2b cells also express and secrete substantial amounts of the anti-inflammatory cytokine IL-10 while producing low levels of IL-12 [31]. In the context of cancer and infectious diseases, M2b macrophages promote tumor progression and facilitate parasitic, bacterial, and fungal infections by attenuating immune and inflammatory responses[32]. M2c exhibits potent anti-inflammatory functions and facilitates efferocytosis by engulfing apoptotic cells, which is induced by glucocorticoids or IL-10-dependent M-CSF signaling. The XCL2 and CXCL16 genes are highly expressed in M2c[33]. Our results demonstrated a mild pro-proliferative effect of M2c macrophages on pulmonary artery smooth muscle cells, which was markedly less pronounced compared to M2a.This also indirectly suggests that M2a macrophages are the primary cells involved in the progression of PAH. M2d macrophages, identified as a novel subset of M2 macrophages and also referred to as tumor-associated macrophages (TAMs), can be induced through co-stimulation with TLR ligands and A2 adenosine receptor (A2R) agonists or interleukin-6 (IL-6) [34]. IL-10 and VEGF signaling in M2d macrophages may represent potential therapeutic targets for mucosal healing[35]. These results indicate that M2a macrophages promote the progression of PAH and reach a relatively stable state in the later stages of the disease.

PTX is a natural tricyclic diterpenoid that inhibits tumor cell proliferation by suppressing both Smad2 expression and Smad3 phosphorylation [36,37]. A recent study by Feng et al. further demonstrated that PTX ameliorates pulmonary vascular remodeling through inhibition of FOXO1-mediated autophagy in smooth muscle cells [38]. In breast tumors, PTX promotes macrophage polarization from the M2 to the M1 phenotype via the TLR4-NF-κB pathway, thereby exerting antitumor effects [39]. Notably, TLR4 (or other TLRs) activates NF-κB or IRF family members in macrophages, driving their polarization toward the M1 phenotype. Following PTX intervention in PAH rats, we found that PTX could not reverse PAH in the early stage but improved pulmonary artery pressure and RVHI in the later stage, while also reducing the accumulation of M2a macrophages around the pulmonary arteries. This suggests that PTX may ameliorate PAH by inhibiting M2a polarization. Additionally, given the high expression of TLR4 and NF-κB proteins in pulmonary tissue macrophages, we confirmed in vitro that PTX suppresses NF-κB gene expression, thereby inhibiting M2a polarization and attenuating its promotive effect on pulmonary artery smooth muscle cell proliferation, ultimately alleviating PAH.

### Conclusions

In this study, we demonstrated the promotive role of M2a macrophages (an M2 subtype) in PAH and confirmed that paclitaxel could inhibit M2a polarization by suppressing the NF-κB pathway. Nevertheless, our research has certain limitations. First, in the cell experiments, we did not perform knockout of M2a-related genes for reverse validation. Second, we did not overexpress M2a genes through plasmid transfection to further confirm their functional role. These aspects will be the focus of our future investigations. Overall, these findings provide a theoretical foundation for developing more precise biologic therapies targeting the M2 subtype for the treatment of PAH.

## ANIMAL ETHICS APPROVAL DECLARATION

All animal experimental and surgical procedures were approved by the Experimental Animal Ethics Committee of West China Second University Hospital of Sichuan University (Ethics number:20211123-068)., and were performed in accordance with the U.S. National Institutes of Health Guide for the Care and Use of Laboratory Animals.

## DATA AVAILABILITY

All supporting information is available from the figshare database (https://doi.org/10.6084/m9.figshare.29885750)

## REFERENCES

1. Kovacs G, Dumitrescu D, Barner A, Greiner S, Grünig E, Hager A, et al. Definition, clinical classification and initial diagnosis of pulmonary hypertension: Updated recommendations from the Cologne Consensus Conference 2018. Int J Cardiol. 2018; 272S:11–19. doi: 10.1016/j.ijcard.2018.08.083.

2. Avitabile CM, Vorhies EE, Ivy DD. Drug Treatment of Pulmonary Hypertension in Children[J]. Paediatr Drugs, 2020; 22(2):123–147. doi: 10.1007/s40272-019-00374-2.

3. Wu Y, Liu H-M, Gu L, Li Q-W, Zhu L. Prostacyclins and pulmonary arterial hypertension in children. Eur Rev Med Pharmacol Sci. 2022; 26(1):37–45. doi: 10.26355/eurrev_202201_27745.

4. Berghausen EM, Feik L, Zierden M, Vantler M, Rosenkranz S. Key inflammatory pathways underlying vascular remodeling in pulmonary hypertension. Herz. 2019; 44(2):130–137. doi: 10.1007/s00059-019-4795-6.

5. Kassa B, Mickael C, Kumar R, Sanders L, Koyanagi D, Saavedra DH, et al. Paclitaxel blocks Th2-mediated TGF-β activation in Schistosoma mansoni-induced pulmonary hypertension. Pulm Circ. 2019;9(1):2045894018820813. doi: 10.1177/2045894018820813.

6. Braithwaite AT, Marriott HM, Lawrie A. Divergent Roles for TRAIL in Lung Diseases. Front Med (Lausanne). 2018: 5:212. doi: 10.3389/fmed.2018.00212.

7. Murray PJ. Macrophage Polarization[J]. Annu Rev Physiol. 2017;79:541–566. doi: 10.1146/annurev-physiol-022516-034339.

8. Vergadi E, Chang MS, Lee C, Liang QD, Liu XL, Gonzalez AF, et al. Early macrophage recruitment and alternative activation are critical for the later development of hypoxia-induced pulmonary hypertension[J]. Circulation, 2011; 123(18):1986–95. doi: 10.1161/CIRCULATIONAHA.110.978627.

9. Sezginer O, Unver N. Dissection of pro-tumoral macrophage subtypes and immunosuppressive cells participating in M2 polarization. Inflamm Res. 2024;73(9):1411–1423. doi: 10.1007/s00011-024-01907-3.

10. Li H, Yang H, Ma B, Qiao J, Chen F, Wang P, et al. Dual-functionalized graphene oxide induces M2a and M2c macrophage polarization to orchestrate inflammation and tissue remodeling. J Mater Chem B. 2025: 13(30):9182–9202. doi: 10.1039/d5tb00474h.

11. Wanderley CW, Colón DF, Luiz JPM, Oliveira FF, Viacava PR, Leite CA, et al. Paclitaxel Reduces Tumor Growth by Reprogramming Tumor-Associated Macrophages to an M1 Profile in a TLR4-Dependent Manner[J]. Cancer Res. 2018; 78(20):5891–5900. doi: 10.1158/0008-5472.CAN-17-3480.

12. Katz MG, Fargnoli AS, Gubara SM, Bisserier M, Sassi Y, Bridges CR, et al. The Left Pneumonectomy Combined with Monocrotaline or Sugen as a Model of Pulmonary Hypertension in Rats. J Vis Exp. 2019: (145):10.3791/59050. doi: 10.3791/59050

13. Sztuka K, Stroschein MJ. Animal models of pulmonary arterial hypertension: A systematic review and meta-analysis of data from 6126 animals. Pharmacol Res. 2017;125(Pt B):201–214. doi: 10.1016/j.phrs.2017.08.003

14. Badesch DB, Raskob GE, Elliott CG, Krichman AM, Farber HW, Frost AE, et al. Pulmonary arterial hypertension: baseline characteristics from the REVEAL registry[J]. Chest. 2010; 137(2):376–387

15. Tobal R, Potjewijd J, van Empel VPM, Ysermans R, Schurgers LJ, Reutelingsperger CP, et al. Vascular Remodeling in Pulmonary Arterial Hypertension: The Potential Involvement of Innate and Adaptive Immunity. Front Med (Lausanne). 2021;:8:806899. doi: 10.3389/fmed.2021.806899.

16. Vaillancourt M, Ruffenach G, Meloche J, Bonnet S. Adaptation and remodelling of the pulmonary circulation in pulmonary hypertension[J]. Can J Cardiol. 2015; 31(4):407–15. doi: 10.1016/j.cjca.2014.10.023.

17. Tian W, Jiang XG, Tamosiuniene R, Sung YK, Qian J, Dhillon, G, et al. Blocking Macrophage Leukotriene B4 Prevents Endothelial Injury and Reverses Pulmonary Hypertension.Sci Transl Med. 2013;5(200):200ra117. doi: 10.1126/scitranslmed.3006674.

18. Qiu H, Zhang Y, Li Z, Jiang P, Guo S, He Y, et al. Donepezil Ameliorates Pulmonary Arterial Hypertension by Inhibiting M2-Macrophage Activation. Front Cardiovasc Med. 2021; 8:639541. doi: 10.3389/fcvm.2021.639541.

19. Tan G, Juan C, Mao Y, Xue G, Fang Z..Inhibition of DLL4/Notch Signaling Pathway Promotes M2 Polarization and Cell Proliferation in Pulmonary Arterial Hypertension. ACS Omega. 2024; 9(36):37923–37933. doi: 10.1021/acsomega.4c04307

20. Fan Y, Hao, YJ, Gao D, Li GT, Zhang ZL. Phenotype and function of macrophage polarization in monocrotaline-induced pulmonary arterial hypertension rat model. Physiol Res. 2021; 70(2):213–226. doi: 10.33549/physiolres.934456

21. Yoshida T, Nagaoka T, Nagata Y, Suzuki Y, Tsutsumi T, Kuriyama S, et al. Periostin-related progression of different types of experimental pulmonary hypertension: A role for M2 macrophage and FGF-2 signalling. Respirology. 2022;27(7):529–538. doi: 10.1111/resp.14249

22. Hourani T., Holden J.A., Li W., Lenzo J.C., Hadjigol S., Brien-Simpson N.MO. Tumor Associated Macrophages: Origin, Recruitment, Phenotypic Diversity, and Targeting. Front. Oncol. 2021;11:788365. doi: 10.3389/fonc.2021.788365.

23. Orecchioni M., Ghosheh Y., Pramod A.B., Ley K. Macrophage Polarization: Different Gene Signatures in M1(LPS+) vs. Classically and M2(LPS-) vs. Alternatively Activated Macrophages. Front. Immunol. 2019; 10:1084. doi: 10.3389/fimmu.2019.01084

24. Biswas S.K., Mantovani A. Macrophage plasticity and interaction with lymphocyte subsets: Cancer as a paradigm. Nat. Immunol. 2010;11:889–896. doi: 10.1038/ni.1937

25. Edwards J.P., Zhang X., Frauwirth K.A., Mosser D.M. Biochemical and functional characterization of three activated macrophage populations. J. Leukoc. Biol. 2006;80:1298–1307. doi: 10.1189/jlb.0406249

26. Little A.C., Pathanjeli P., Wu Z., Bao L., Goo L.E., Yates J.A., et al. IL-4/IL-13 Stimulated Macrophages Enhance Breast Cancer Invasion Via Rho-GTPase Regulation of Synergistic VEGF/CCL-18 Signaling. Front. Oncol. 2019;9:456. doi: 10.3389/fonc.2019.00456

27. Zhang Q, Sioud M. Tumor-Associated Macrophage Subsets: Shaping Polarization and Targeting..Int J Mol Sci. 2023; 24(8):7493. doi: 10.3390/ijms24087493.PMID: 37108657

28. Stein M., Keshav S., Harris N., Gordon S. Interleukin 4 potently enhances murine macrophage mannose receptor activity: A marker of alternative immunologic macrophage activation. J. Exp. Med. 1992; 176:287–292. doi: 10.1084/jem.176.1.287

29. Huang X, Li Y, Fu MG, Xin HB. Activating THP1-derived macrophage in vitro. Methods Mol Biol. 2018;1784:119–126. doi: 10.1007/978-1-4939-7837-3

30. Mosser DM, Edwards JP. Exploring the full spectrum of macrophage activation. Nat Rev Immunol. 2008;8(12):958–69. doi: 10.1038/nri2448.

31. Yue Y, Yang X, Feng K, Wang LX, Hou J, Mei B, et al. M2b macrophages reduce early reperfusion injury after myocardial ischemia in mice: a predominant role of inhibiting apoptosis via A20. Int J Cardiol. 2017; 245:228–235. doi: 10.1016/j.ijcard.2017.07.085.

32. Asai A, Tsuchimoto Y, Ohama H, Fukunishi S, Tsuda Y, Kobayashi M, et al. Host antitumor resistance improved by the macrophage polarization in a chimera model of patients with HCC. Oncoimmunology. 2017;6(4):e1299301. doi: 10.1080/2162402X.2017.1299301.

33. Lai YS, Putra RBDS, Aui S-P, Chang KT. M2C polarization by Baicalin enhances efferocytosis via Upregulation of MERTK receptor. Am J Chin Med. 2018; 46(8):1899–1914. doi: 10.1142/S0192415X18500957.

34. Zhang Q, Sioud M. Tumor-Associated Macrophage Subsets: Shaping Polarization and Targeting. Int J Mol Sci. 2023; 24(8):7493. doi: 10.3390/ijms24087493.

35. Lai W, Xian C, Chen M, Luo D, Zheng J, Zhao S, et al. Single-cell and bulk transcriptomics reveals M2d macrophages as a potential therapeutic strategy for mucosal healing in ulcerative colitis. Int Immunopharmacol. 2023; 121:110509. doi: 10.1016/j.intimp.2023.110509

36. Cheng H, Chen T, Tor M, Park D, Zhou QY, Huang JB et al. A High-Throughput Screening Platform Targeting PDLIM5 for Pulmonary Hypertension [J]. J Biomol Screen. 2016; 21(4): 333–41. doi: 10.1177/1087057115625924

37. Yin Y, Wu X, Yang Z, Zhao J, Wang XS, Zhang QY, et al. The potential efficacy of R8-modified paclitaxel-loaded liposomes on pulmonary arterial hypertension[J]. Pharm Res. 2013; 30(8):2050–2062. doi: 10.1007/s11095-013-1058-8.

38. Feng W, Wang J, Yan X, Zhai C, Shi WH, Wang QT, et al. Paclitaxel alleviates monocrotaline-induced pulmonary arterial hypertension via inhibition of FoxO1-mediated autophagy [J]. Naunyn Schmiedebergs Arch Pharmacol. 2019; 392(5): 605–613. doi: 10.1007/s00210-019-01615-4

39. Wanderley CW, Colón DF, Luiz JPM, Oliveira FF, Viacava PR, Leite CA, et al. Paclitaxel Reduces Tumor Growth by Reprogramming Tumor-Associated Macrophages to an M1 Profile in a TLR4-Dependent Manner[J]. Cancer Res. 2018;78(20):5891–5900. doi: 10.1158/0008-5472.CAN-17-3480

